# Using Domain-Knowledge to Assist Lead Discovery in Early-Stage Drug Design

**DOI:** 10.1101/2021.07.09.451519

**Authors:** Tirtharaj Dash, Ashwin Srinivasan, Lovekesh Vig, Arijit Roy

## Abstract

We are interested in generating new small molecules which could act as inhibitors of a biological target, when there is limited prior information on target-specific inhibitors. This form of drug-design is assuming increasing importance with the advent of new disease threats for which known chemicals only provide limited information about target inhibition. In this paper, we propose the combined use of deep neural networks and Inductive Logic Programming (ILP) that allows the use of symbolic domain-knowledge (*B*) to explore the large space of possible molecules. Assuming molecules and their activities to be instances of random variables *X* and *Y*, the problem is to draw instances from the conditional distribution of *X*, given *Y, B* (*D_X|Y,B_*). We decompose this into the constituent parts of obtaining the distributions *D_X|B_* and *D_Y|X,B_*, and describe the design and implementation of models to approximate the distributions. The design consists of generators (to approximate *D_X|B_* and *D_X|Y,B_*) and a discriminator (to approximate *D_Y|X,B_*). We investigate our approach using the well-studied problem of inhibitors for the Janus kinase (JAK) class of proteins. We assume first that if no data on inhibitors are available for a target protein (JAK2), but a small numbers of inhibitors are known for homologous proteins (JAK1, JAK3 and TYK2). We show that the inclusion of relational domain-knowledge results in a potentially more effective generator of inhibitors than simple random sampling from the space of molecules or a generator without access to symbolic relations. The results suggest a way of combining symbolic domain-knowledge and deep generative models to constrain the exploration of the chemical space of molecules, when there is limited information on target-inhibitors. We also show how samples from the conditional generator can be used to identify potentially novel target inhibitors.

## 1 Introduction

Co-opting Hobbes, the development of a new drug is difficult, wasteful, costly, uncertain, and long. AI techniques have been trying to change this [1], especially in the early stages culminating in “lead discovery”. Figure 1 shows the steps involved in this stage of drug-design. In the figure, library screening can be either done by actual laboratory experiments (high-throughput screening) or computationally (virtual screening). This usually results in many false-positives. Hit Confirmation refers to additional assays designed to reduce false-positives. QSAR (quantitative- or qualitative structure-activity relations) consists of models for predicting biological activity using physico-chemical properties of hits. The results of prediction can result in additional confirmatory assays for hits, and finally in one or more “lead” compounds that are taken forward for pre-clinical testing. This paper focuses on the problem of lead discovery that goes beyond the efficient identification of chemicals within the almost unlimited space of potential molecules. This space has been approximately estimated at about 10^60^ molecules. A very small fraction of these have been synthesised in research laboratories and by pharmaceutical companies. An even smaller number are available publicly: the well-known ChEMBL database [2] of drug-like chemicals consists of about 10^6^ molecules. Any early-stage drug-discovery pipeline that restricts itself to in-house chemicals will clearly be self-limiting. This is especially the case if the leads sought are for targets in new diseases, for which very few “hits” may result from existing chemical libraries. While a complete (but not exhaustive) exploration of the space of 10^60^ molecules may continue to be elusive, we would nevertheless like to develop an effective way of sampling from this space.

**Fig. 1.**
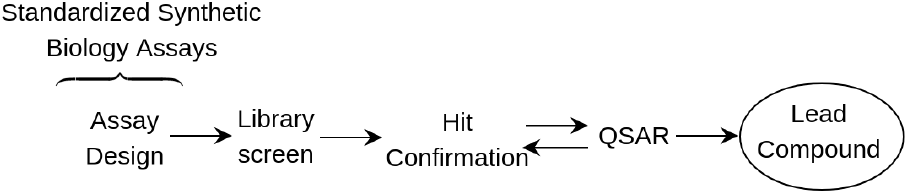
Early-stage drug-design (adapted from [3]).

We would like to implement the QSAR module as a generator of new molecules, conditioned on the information provided by the hit assays, and on domain-knowledge. Our position is that inclusion of domain-knowledge allows the development of more effective conditional distributions than is possible using just the hit assays. Figure 2 is a diagrammatic representation of an ideal conditional generator of the kind we require. The difficulty of course is that none of the underlying distributions are known. In this paper, we describe a neural-symbolic implementation to construct approximations for the distributions.

**Fig. 2.**
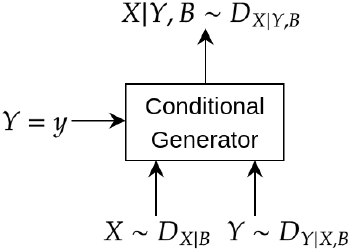
An ideal conditional generator for instances of a random-variable denoting data (*X*) given a value for a random-variable denoting labels (*Y*) and domain-knowledge (*B*). Here, *Z* ~ *D* denotes a random variable *Z* is distributed according to the distribution *D*. If the distributions shown are known, then a a value for *X* is obtainable through the use of Bayes rule, either exactly or through some form approximate inference.

## 2 System Design and Implementation

We implement an approximation to the ideal conditional generator using a generator-discriminator combination (see Fig. 3). We have decomposed the domain-knowledge *B* in Fig. 2 into constraints relevant just to the molecule-generator *B_G_* and the knowledge relevant to the prediction of activity *B_D_* (that is, *B* = *B_G_*∪*B_D_* and *P* (*X*|*B*) = *P* (*X*|*B_G_*) and *P* (*Y*|*X, B*) = *P* (*Y*|*X, B_D_*)). The discriminator module approximates the conditional distribution *D_Y|X,B_*, and the combination of the unconditional generator and filter approximates the distribution *D_X|B_*. The conditional generator then constructs an approximation to *D_X|Y,B_*. For the present, we assume the unconditional generator and discriminator are pre-trained: details will be provided below. The discriminator is a BotGNN [4]. This is a form of graph-based neural network (GNN) that uses graph-encodings of most-specific clauses (see [5]) constructed using symbolic domain-knowledge *B_D_*.

**Fig. 3.**
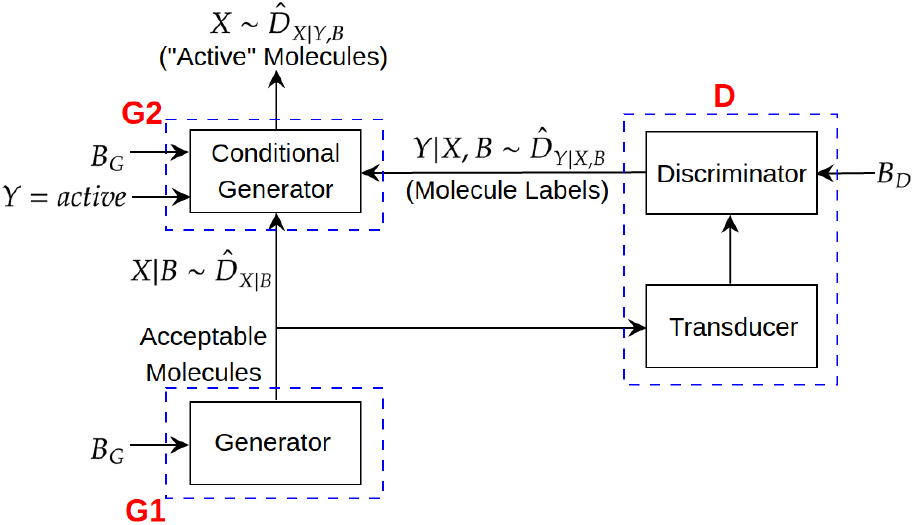
Training a conditional generator for generating “active” molecules. For the present, we assume the generator (G1) and discriminator (D) have already been trained (the G1 and D modules generate acceptable molecules and their labels respectively: the 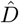’s are approximations to the corresponding true distribution). The Transducer converts the output of G1 into a form suitable for the discriminator. Actual implementations used in the paper will be described below.

The generator-discriminator combination in Fig. 3 constitutes the QSAR module in Fig. 1. An initial set of hits is used to train the discriminator. The conditional generator is trained using the initial set of hits and the filtered samples from the unconditional generator and the labels from the discriminator. Although out of the scope of this paper, any novel molecules generated could then be synthesised, subject to hit confirmation, and the process repeated.

### Generating Acceptable Molecules

The intent of module G1 is to produce an approximation to drawing samples (in our case, molecules) from 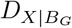. We describe the actual *B_G_* used for experiments in Sec. 3.1: for the present it is sufficient to assume that for any instance *X* = *x*, if *B_G_* ∧ *X* = *x* |= ◻ then *Pr*(*x|B_G_*) = 0. Here, we implement this by a simple rejection-sampler that first draws from some distribution over molecules and rejects the instances that are inconsistent with *B_G_*.

For drawing samples of molecules, we adopt the text-generation model proposed in [6]. Our model takes SMILES representations of molecules as inputs and estimates a probability distribution over these SMILES representations. Samples of molecules are then SMILES strings drawn from this distribution.

The SMILES generation module is shown in Fig. 4. The distribution of molecules (SMILES strings) is estimated using a variational autoencoder (VAE) model. The VAE model consists of an encoder and a decoder, both based on LSTM-based RNNs [7]. This architecture forms a SMILES encoder with the Gaussian prior acting as a regulariser on the latent representation. The decoder is a special RNN model that is conditioned on the latent representation.

**Fig. 4.**
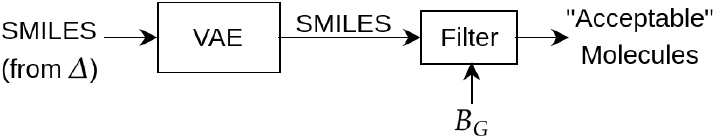
Training a generator for acceptable molecules. Training data consists of molecules, represented as SMILES strings, drawn from a database *Δ*. The VAE is a model constructed using the training data and generates molecules represented by SMILES strings. *B_G_* denotes domain-knowledge consisting of constraints on acceptable molecules. The filter acts as a rejection-sampler: only molecules consistent with *B_G_* pass through.

The architecture of the VAE model is shown in Fig. 5. The SMILES encoding involves three primary modules: (a) embedding module: constructs an embedding for the input SMILE; (b) highway module: constructs a gated information-flow module based on highway network [8]; (c) LSTM module: responsible for dealing with sequence. The modules (b) and (c) together form the encoder module. The parameters of the Gaussian distribution is learnt via two fully-connected networks, one each for ***μ*** and ***σ***, which are standard sub-structures involved in a VAE model. The decoder module (or the generator) consists of LSTM layers followed by a fully-connected (FC) layer. We defer the details on architecture-specific hyperparameters to Sec. 3.2. The loss function used for training our VAE model is a weighted version of the reconstruction loss and KL-divergence between VAE-constructed distribution and the Gaussian prior 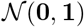.

**Fig. 5.**
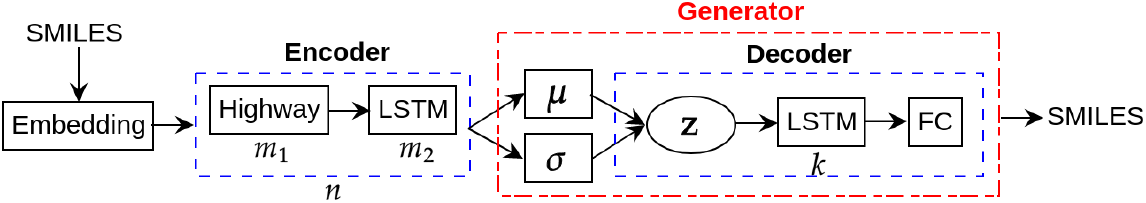
Architecture of the VAE in Fig. 4. *m*_1,2_, *n, k* represent the number of blocks.

### Obtaining Labels for Acceptable Molecules

The intent of module D is to produce an approximation to drawing samples (in our case, labels for molecules) from 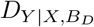. We describe the actual *B_D_* used for experiments in Sec. 3.1. The discriminator in D is a BotGNN [4], which is a form of graph neural network (GNN) constructed from data (as graphs) and background knowledge (as symbolic relations or propositions) using mode-directed inverse entailment (MDIE [5]). In this work, data consists of graph-based representations of molecules (atoms and bonds), and *B_D_* consists of symbolic domain-relations applicable to the molecules. The goal of the discriminator is to learn a distribution over class-labels for any given molecules. Fig. 6 shows the block diagram of the discriminator block.

**Fig. 6.**
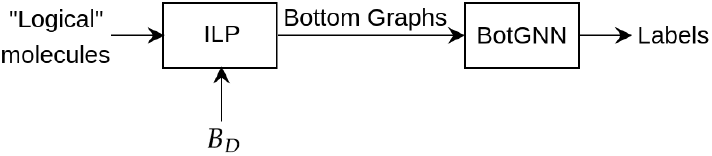
Discriminator based on BotGNN. “Logical” molecules refers to a logic-based representation of molecules. Bottom-graphs are a graph-based representation of most-specific (“bottom”) clauses constructed for the molecules by an ILP implementation based on mode-directed inverse entailment.

### Generating Active Molecules

The intent of module G2 is to produce an approximation to drawing from *D_X|Y,B_*. That is, we want to draw samples of molecules, given a label for the molecule and domain-knowledge *B*. We adopt the same architecture as the generator used for drawing from 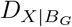 above, with a simple modification to the way the SMILES strings are provided as inputs to the model. We prefix each SMILES string with a class-label: *y* = 1 or *y* = 0 based on whether the molecule is an active or inactive inhibitor, respectively. The VAE model is also able to accommodate any data that may already be present about the target, or about related targets (it is assumed that such data will be in the form of labelled SMILES strings).

## 3 System Testing

Our aim is to perform a controlled experiment to assess the effect on system performance of the inclusion of high-level symbolic domain-knowledge. Specifically: we investigate the effect on the generation of new inhibitors for the target when: (a) No domain-knowledge is available in the form of symbolic relations (but some knowledge is available in a propositional form); and (b) Some domain-knowledge is available in form of symbolic relations. We intend to test if the system is able to generate possible new inhibitors in case (a); and if the performance of the system improves in case (b).

### 3.1 Materials

#### Data

The data used are as follows. (a) ChEMBL dataset [2]: 1.9 million molecules; used to train the generator for legal molecules (G1); (b) JAK2 [9]: 4100 molecules (3700 active); used to test the conditional generator (G1) and to build the proxy model for hit confirmation (see Method section below); (c) JAK2 Homologues (JAK1, JAK3 and TYK2) [9]: 4300 molecules (3700 active); used to train the discriminator (D) and train the conditional generator (G2).

#### Domain-Knowledge

We use the following categories of domain-knowledge (also see Appendix A). (a) Molecular Constraints [9,10]: Logical constraints on acceptable molecules, including standard validity checks (based on molecular properties); (b) Molecular Properties [9]: Bulk-properties of molecules (propositional in nature); (c) Molecular Relations [11]: Logical statements defining ring-structures and functional groups (relational in nature).

#### Algorithms and Machines

We use the following software. (a) RDKit [10]: Molecular modelling software used to compute molecular properties and check for the validity of molecules; (b) Chemprop [12]: Molecular modelling software used to build a proxy model for hit confirmation; (c) Transducer: In-house software to convert representation from SMILES to logic; (d) Aleph [13]: ILP engine used to generate most-specific clauses for BotGNN; (e) BotGNN [4]: Discriminator for acceptable molecules capable of using relational and propositional domain knowledge; (f) VAE [14]: Generative deep network used for generators. We used PyTorch for the implementation of BotGNN and VAE models, and Aleph was used with YAP.

Our experimental works were distributed across two machines: (a) The discriminator (D) was built on a Dell workstation with 64GB of main memory, 16-core Intel Xeon 3.10GHz processors, an 8GB NVIDIA P4000 graphics processor; (b) The generators (G1, G2) are built on an NVIDIA-DGX1 station with 32GB Tesla V100 GPUs, 512GB main memory, 80-core Intel Xeon 2.20GHz processors.

### 3.2 Methods

We describe the procedure adopted for a controlled experiment comparing system performance in generating potential inhibitors when: (a) domain-knowledge is restricted to commonly used bulk-properties about the molecules; and (b) domain-knowledge includes information about higher-level symbolic relations consisting of ring-structures and functional groups, along with the information in (a). In either case, the method used to generate acceptable molecules (from module G1 in Fig. 3) is the same.

Let *B*_0_ denote domain-knowledge consisting of bulk-molecular properties used in the construction of QSARs for novel inhibitors; *B*_1_ denote the the definitions in *B*_0_ along with first-order relations defining ring-structures and functional-groups used in the construction of QSAR relations; and *B_G_* denote the domain-knowledge consisting of constraints on acceptable molecules (see “Domain-Knowledge” in Sec. 3.1). Let *Tr* denote the data available on inhibitors for JAK1, JAK3 and TYK2; and *Te* denote the data available on inhibitors for JAK2 (see “Data” in Sec. 3.1). Let *Δ* denote a database of (known) legal molecules. Then:

1. Construct a generator for possible molecules given *Δ* (the generator in module G1 of Fig. 3).
2. For *i* = 0, 1
  a. Let 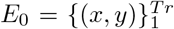, where *x* is a molecule in *Tr* and *y* is the activity label obtained based on a threshold *θ* on the minimum activity for active inhibition.
  b. Let *B_D_* = *B_i_*
  c. Construct a discriminator (for module D in Fig. 3) using *E*_0_ and the domain-knowledge *B_D_* (see Sec. 2);
  d. Sample a set of possible molecules, denoted as *N*, from the generator constructed in Step 1;. Let *N*′ ⊆ *N* be the set of molecules found to be acceptable given the constraints in *B_G_* (that is, *N*′ is a sample from 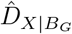);
  e. For each acceptable molecule *x* obtained in Step 2d above, let *y* be the label with the highest probability from the distribution 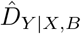 constructed by the discriminator in Step 2c. Let 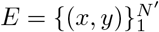
  f. Construct the generator model (for module G2 in Fig. 3) using *E*_0_ ∪ *E*.
  g. Sample a set of molecules, denoted as *M_i_*, from the generator in Step 2f;
  h. Let 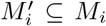 be the molecules found to be acceptable given the constraints in *B_G_* (that is, 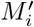 is a sample from 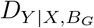)
3. Assess the samples *M*_0,1_ obtained in Step 2g above for possible new inhibitors of the target, using the information in *Te*

The following details are relevant:

– For experiments here *Δ* is the ChEMBL database, consisting of approximately 1.9 million molecules. The generator also includes legality checks performed by the RDKit package, as described in Sec. 2;
– Following [9], *θ* = 6.0. That is, all molecules with pIC50 value ≥ 6.0 are taken as “active” inhibitors;
– The discriminator in Step 2c is a BotGNN. We follow the procedure and parameters described in [4] to construct BotGNN. We use GraphSAGE [15] for the convolution block in the GNN. This is based on the results shown in [4] for including symbolic domain knowledge for graph-based data (like molecules);
– The generators in Steps 1 and 2f are based on the VAE model described earlier. The hyperparameters are as follows: vocabulary length is 100, embedding-dimension is 300, number of highway layers is 2, number of LSTM layers in the encoder is 1 with hidden size 512, and the type is bidirectional, number of LSTM layers in the decoder is 2, each with hidden size 512, dimension of latent representation (**z**) is 100.
– To make our generator robust to noise and to be generalised, we also use a word-dropout technique. This technique is identical to the standard practice of dropout in deep learning except that here the tokens to the decoder are replaced by ‘*unknown*’ tokens with certain probabilities. Here we call it the word-dropout rate and fix it at 0.5.
– The reconstruction loss coefficient is 7. We use cost-annealing [6] for the KLD-coefficient during training. We use the Adam optimizer [16] with learning rate of 0.0001; training batch-size is 256.
– In Step 2d, |*N* | = 30, 000. The *B_G_* provided here results in |*N*′| = 18, 000;
– In Step 2g, |*M*_0_| = |*M*_1_| = 5000.
– The acceptable molecules 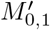 after testing for consistency with *B_G_* are assessed along two dimensions:
  a. *Activity* : In the pipeline described in Fig. 1 assessment of activity would be done by *in vitro* by hit confirmation assays. Here we use a proxy assessment for the result of the assays by using an *in silico* predictor of pIC50 values constructed from the data in *Te* on JAK2 inhibitors. The proxy model is constructed by a state-of-the-art activity prediction package (Chemprop [12]: details of this are in the Appendix).^4^ We are interested in comparing the proportions of generated molecules predicted as “active”;
  b. *Similarity* : we want to assess how similar the molecules generated are to the set of active JAK2 inhibitors in *Te*.^5^ A widely used measure for this is the Tanimoto (Jacquard) similarity: molecules with Tanimoto similarity > 0.75 are usually taken to be similar. We are interested in the proportion of molecules generated that are similar to known target inhibitors in *Te*;
  Each sample of molecules *M_i_* drawn from the conditional generator can therefore be represented by a pair (*a_i_, b_i_*) denoting the values of the proportions in (a) and (b), and (c) above. We will call this pair the “performance summary” of the set *M_i_*;
– We compare performance summaries of sets of molecules in two ways. First, a performance summary *P_i_* = (*a_i_, b_i_*) can be compared against the performance summary *P_j_* = (*a_j_, b_j_*) in the obvious lexicographic manner. That is, *P_i_* is better than *P_j_* if [(*a_i_ > a_j_*)] or [(*a_i_* = *a_j_*) ∧ (*b_i_ > b_j_*)]. Secondly, since all the elements of a performance summary are proportions, we are able to assess if the differences in corresponding values are statistically significant. This is done using a straightforward hypothesis test on proportions. Given an estimate *p* of a proportion of *N* instances, the distribution of proportions is approximately Normal, with mean *p* and s.d. 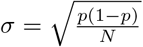. For testing the hypothesis *p_j_ < p_i_* at a 95% confidence level the critical value from tables of the standard normal distribution is 1.65. That is, if *p_j_*< 1.65*σ* we will say the difference is statistically significant at the 95% level of confidence.

### 3.3 Results

A summary of the main results obtained is in Fig. 7. The principal points in this tabulation are these: (1) The performance of the system with *B_D_* = *B*_1_ is better than with *B_D_* = *B*_0_ or simple random draw of molecules; and (2) The differences in proportions for Activity and Similarity are statistically significant at the 95% confidence level. Taken together, these results suggest that the inclusion of symbolic relations can make a significant difference to the performance of the generation of active molecules. We turn next to some questions of relevance to these results:

**Fig. 7.**
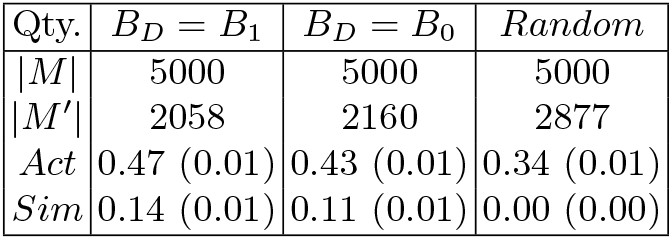
Summary of system performance. *B_D_* = *B*_1_ denotes that the discriminator has access to both propositional and relational domain-knowledge; *B_D_* = *B*_0_ denotes that the discriminator has access to propositional domain-knowledge only. *Random* denotes a random draw of molecules from the unconditional molecule generator G1. *M* denotes the set of molecules drawn (from the conditional generator, or from the unconditional generator for *Random*). *M*′ denotes the set of acceptable molecules generated in the sample of *M* molecules (acceptable molecules satisfy molecular constraints defined on molecular properties). *Act* denotes the proportion of *M*′ that are predicted active (the proxy model predicts an pIC50 ≥ 6.0); *Sim* denotes the proportion of *M*′ that are similar to active target inhibitors (Tanimoto similarity to active JAK2 inhibitors > 0.75). The numbers in parentheses denote the standard deviation in the corresponding estimate.

#### Better Discriminators?

A question arises on whether the differences in proportions would be different if we had compared against a different discriminator capable of using *B_D_* = *B*_0_. Since *B*_0_ is essentially propositional in nature, any of the usual statistical discriminative approaches could be used. We have found replacing the BotGNN with an MLP with hyper-parameter tuning resulted in significantly worse performance than a BotGNN with *B_D_* = *B*_1_. We conjecture that similar results will be obtained with other kinds of statistical models. On the question of whether better discriminators are possible for *B_D_* = *B*_1_, we note results in [4] show BotGNNs performance to be better than techniques based on propositionalisation or a direct use of ILP. Nevertheless, better BotGNN models than the one used here may be possible. For example, we could construct an activity prediction model for the JAK2 homologues using a state-of-the-art predictor like Chemprop. The prediction of this model could be used as an additional molecular property by the BotGNN.

#### Better Generators?

Our generators are simple language models based on variational auto-encoders. Substantial improvements in generative language models (for example, the sequence models based on attention mechanism [17,18]) suggest that the generator could be much better. In addition, the rejection-sampling approach we use to discard sample instances that fail constraints in *B_G_* is inherently inefficient, and we suggest that the results here should be treated as a baseline. The modular design of our system-design should allow relatively easy testing of alternatives.

Related to the question of discriminators is the role of ILP in this work. ILP is used to include domain-knowledge in the construction of the BotGNN discriminator. How important was this use of ILP? A quantitative answer is difficult, but we are able to provide indirect, qualitative evidence for the utility of ILP by comparison against a recent result on the same data in [9]. That work differs from the one here in the following ways: (a) No symbolic domain-knowledge is used in the discrimination step; and (b) Substantially more computation is involved in developing the final generator–the equivalent of module G2 here–through the use of reinforcement learning (RL). The principal concern in [9] is to generate molecules similar to the active inhibitors for JAK2, and the approach results in 5% of the sampled molecules being similar. The corresponding results here are significantly higher: 14% (with *B_D_* = *B*_1_) and 11% (*B_D_* = *B*_0_). Both results were obtained with BotGNNs, without requiring the additional episodic training characteristic of RL. Therefore, we believe BotGNNs have played an important role, both in prediction and in easing computation. Since ILP is necessary for the construction of a BotGNN, their importance to the current system-design follows.^6^

Finally, we consider how samples from the conditional generator can be used to identify potential molecules for synthesis and testing in hit-confirmation assays. We propose a selection based on a combination of (predicted) activity and similarity to the existing inhibitors (when these are unavailable, we would have to rely on models constructed with the target’s homologues). Using these measures, there are two surprising subsets of molecules. Molecules in *S* are those that are similar to JAK2 inhibitors (Tanimoto similarity > 0.75), but have a low predicted activity (substantially lower than 6.0); and molecules in 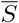 are significantly different to the JAK2 inhibitors (Tanimoto similarity < 0.5), but have a high predicted activity (substantially higher that 6.0).^7^ For the sample in this paper, *S* = Ø. However, 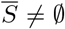 and can provide interesting candidates for novel inhibitors. We exemplify this with a chemical assessment of 3 elements from 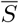. This is shown in Fig. 8. Molecule 1562 is identified as a possible candidate for synthesis and hit confirmation.

**Fig. 8.**
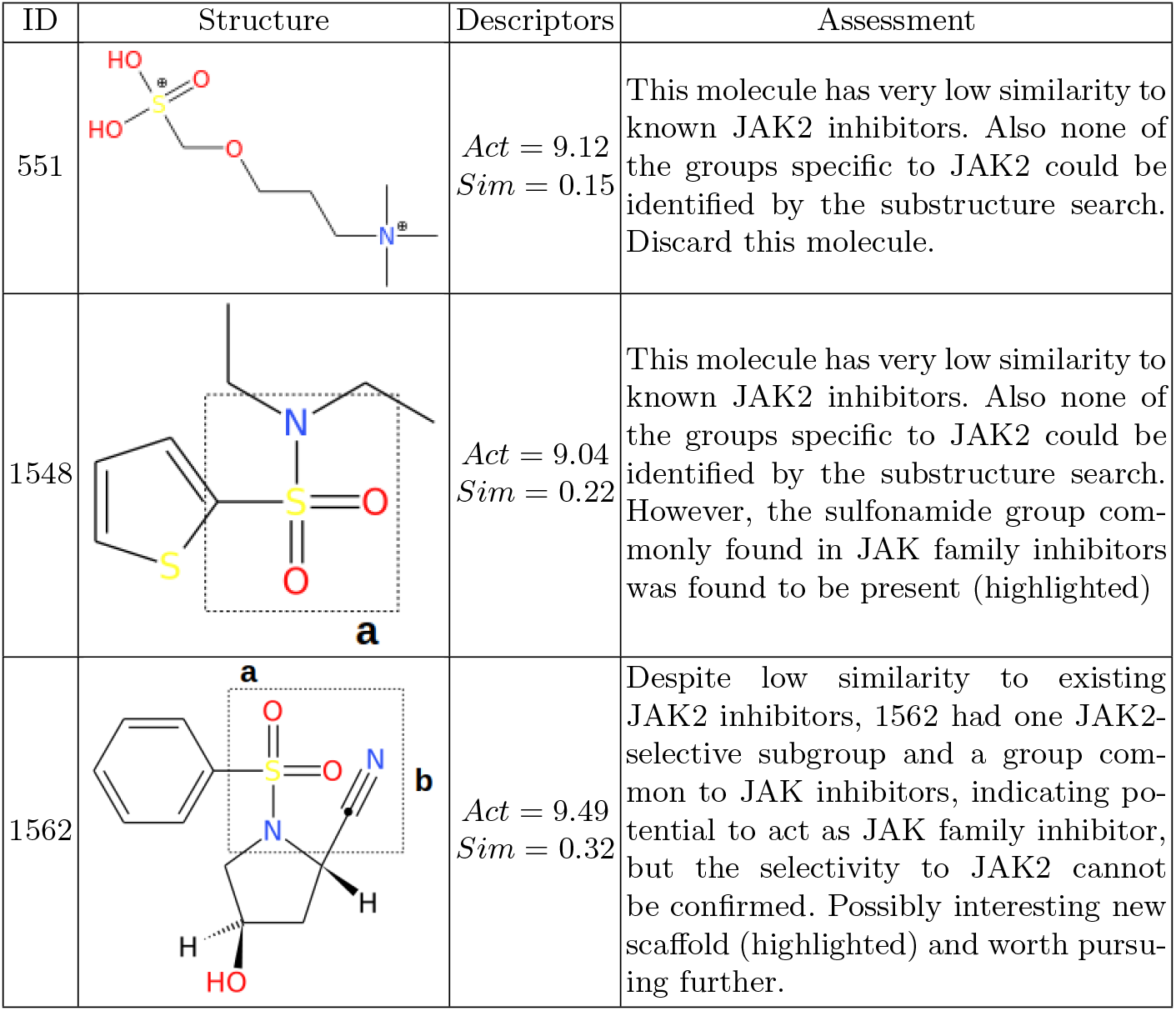
A chemical assessment of possible new JAK2 inhibitors. The molecules are from the sample of molecules from the conditional generator, that are predicted to have high JAK2 activity, and are significantly dissimilar to known inhibitors. The assessment is done by one of the authors (AR), who is a computational chemist. The assessment uses structural features and functional groups identified for the JAK2 site in the literature [9,19,20].

## 4 Related Work

Recent applications of AI-based methods have shown promise in transforming otherwise long and expensive drug discovery process [21,22]. The initial studies were focused on exploring vast yet unexplored chemical space for a better screening library. In [23], a recurrent neural network (RNN) based generative model was trained with a large set of molecules and then fine-tuned with small sets of molecules, which are known to be active against the target. Some other works focus on drug-like property optimization, which helped in biasing the models to generate molecules with specific biological or physical properties of interest. Deep reinforcement learning has been very effective in constructing generative models that could generate novel molecules with the target properties [22,24,25]. The efficiency of these kinds of models to generate chemically valid molecules with optimized properties has improved significantly [9,24]. There are also attempts to build molecule generation models against novel target proteins, where there is a limited ligand dataset for training the model [26].

Recurrent Neural Networks (RNNs) are a popular choice for molecule generation. For example, [27] propose a bidirectional generative RNN, that learns SMILES strings in both directions allowing it to better approximate the data distribution. Attention-based sequence models such as transformers have recently been used for protein-specific molecule generation [28]. There are also generative models, for instance, masked graph modelling in [29], that attempts to learn a distribution over molecular graphs allowing it to generate novel molecule without requiring to dealing with sequences. Some generative modelling techniques for molecule generation are surveyed in [30].

Incorporating domain-knowledge into deep neural networks have shown considerable success over the years. There are several categories of domain knowledge that has been incorporated into learning, primarily referring to the way the knowledge is represented [31]. We present here a brief overview of methods that deal with domain-knowledge represented in a relational form. Possibly the earliest approach to integrating this kind of domain-knowledge is propositionalisation [32,33]. It is a technique to transform a relational representation into a propositional single-table representation where each column in the table corresponds a feature that represents a relation constructed from data and domain-knowledge. Propositionalisation is the core technique in construction of deep relational machines [34,35,36]: these are multi-layered perceptrons constructed from propositionalised representation of relational data and domain-knowledge. Recent studies on domain-knowledge inclusion include construction of graph neural networks (GNNs) that can learn not only from relational (graph-structured) data but also symbolic domain-knowledge. For instance, the vertex-enrichment approach in [37] constructs an enriched vertex-labelling for graph-structured data instances by treating available domain relations as hyperedges. Another approach transforms the most-specific (bottom) clauses in ILP into a bipartite graph structures [4]. These graphs are called bottom-graphs. A GNN can be learned from these bottom-graphs, thereby allowing a principled way of integrating symbolic domain-knowledge into GNNs. A recent survey presents a more elaborate discussion on various kinds of domain-knowledge and the methods of their inclusion into deep neural networks [38].

## 5 Concluding Remarks

Incorporating some form of domain-knowledge into AI-based scientific discovery has been emphasised strongly in [39]. A cutting-edge example of this form of scientific discovery is the Robot Scientist [3], the latest generation of which– Eve–is concerned with automating early-stage drug-design. At the heart of Eve is the development of QSAR models. To the best of our knowledge, generation of molecules is restricted to a library of known chemicals; and the use of domain-knowledge is limited to pre-defined features. In this work, we have proposed an approach that can generate novel molecules drawn from the very large space of all possible small molecules, rather than pre-defined libraries; and we use a method that allows the inclusion of relational domain-knowledge. The paper makes the following contributions: (1) We have constructed a complete end-to-end neural-symbolic system that is capable of generating active molecules that may not be in any existing database; (2) We have demonstrated usage of the system on the classic chemical problem on Janus kinase inhibitors. Importantly, working with a computational chemist, we have shown how the system can be used to discover an active molecule based on entirely new scaffolds; (3) The results reaffirm the conclusions from [4] that inclusion of relational domain knowledge through the use of ILP techniques can significantly improve the performance of deep neural networks. To the best of our knowledge, the system-design is the first-of-a-kind combination of neural generative models, techniques from Inductive Logic Programming and symbolic domain-knowledge representation for lead-discovery in early stage drug-design, and is of relevance to platforms like Eve.

Our system design is intentionally modular, to allow “plug-and-play” of discriminators and generators. Indeed, there is already evidence from the construction of language-models that the VAE-based generators we have used could be replaced by transformer-based deep networks. Thus, an immediate next step would be to replace the existing generators with pre-trained language models like GPT-2. We would also expect that molecular constraints would include both hard- and soft-constraints (unlike here, where only hard-constraints are used). This may presage a move to a probabilistic logic representation of the domain-knowledge. On discriminators, BotGNNs continue to be a good choice for inclusion of symbolic knowledge into deep networks, although, as we have pointed out, the BotGNN model could be improved by inclusion as part of domain-knowledge, results from models constructed by programs like like Chemprop (the extensive use of fingerprints by such programs is essentially a form of relational information), and also the possibility of inclusion of 3-dimensional constraints (see for example, [40]). Looking beyond the goal of novel molecule generation, a promising line of research concerns the development of schedules for synthesis of new molecules. Of special interest is to consider if techniques for experiment-selection could be adopted for prioritising molecules for synthesis.

## Acknowledgements

AS is a Visiting Professorial Fellow at UNSW, Sydney; and a TCS Affiliate Professor. We thank Indrajit Bhattacharya for thoughtful discussions on system-design.

## A Domain-Knowledge used in Experiments

The domain constraints in *B_G_* are in the form of constraints on acceptable molecules. These constraints are broadly of two kinds: (i) Those concerned with the validity of a generated SMILES string. This involves various syntax-level checks, and is done here by the RDKit molecular modelling package; (ii) Problem-specific constraints on some bulk-properties of the molecule. These are: molecular weight is in the range (200, 700), the octanol-water partition coefficients (logP) must be below 6.0, and the synthetic accessibility score (SAS) must be below 5.0. We use the scoring approach proposed in [41] to compute the SAS of a molecule based on its SMILES representation.

The domain-knowledge in *B_D_* broadly divides into two kinds: (i) Propositional, consisting of molecular properties. These are: molecular weight, logP, SAS, number of hydrogen bond donors (HBD), number of hydrogen bond acceptor (HBA), number of rotatable bonds (NRB), number of aromatic rings (NumRings), Topological Polar Surface Area (TPSA), and quantitative estimation of drug-likeness (QED); (ii) Relational, which is a collection of logic programs (written in Prolog) defining almost 100 relations for various functional groups (such as amide, amine, ether, etc.) and various ring structures (such as aromatic, non-aromatic, etc.). The initial version of these background relations was used within DMax chemistry assistant [11]. More details on this background knowledge can be found in [4,37].

## B Proxy Model for Predicting Hit Confirmation

A proxy for the results of hit confirmation assays is constructed using the assay results available for the target. This allows us to approximate the results of such assays on molecules for which experimental activity is not available. Of course, such a model is only possible within the controlled experimental design we have adopted, in which information on target inhibition is deliberately not used when constructing the discriminator in D and generator in G2. In practice, if such target-inhibition information is not available, then a proxy model would have to be constructed by other means (for example, using the activity of inhibitors of homologues).

We use the state-of-the-art chemical activity prediction package Chemprop.^8^ We train a Chemprop model using the data consisting of JAK2 inhibitors and their pIC50 values. The parameter settings used are: `class-balance` = TRUE, and `epochs` = 100 (all other parameters were set to their default values within Chemprop). Chemprop partitions the data into 80% for training, 10% validation and 10% for test. Chemprop allows the construction of both classification and regression models. The performance of both kinds of models are tabulated below:

| Partition | Classification<br>(AUC) | Regression<br>(RMSE) |
| --- | --- | --- |
| Valid | 0.9472 | 0.6515 |
| Test | 0.8972 | 0.6424 |

The classification model is more robust, since pIC50 values are on a log-scale. We use the classification model for obtaining the results in Fig. 7, and we use the prediction of pIC50 values from the regression model as a proxy for the results of the hit-confirmation assays.

## Footnotes

4 Such a model is only possible in the controlled experiment here. In practice, no inhibitors would be available for the target and activity values would have to be obtained by hit assays, or perhaps *in silico* docking calculations.

5 Again, this is feasible in the controlled experiment here. In practice, we will have no inhibitors for the target, and we will have to perform this assessment on the data available for the target’s homologues (*T r*).

6 Could we have directly used ILP for constructing the discriminator? Yes, but there is substantial evidence to suggest that the use of ILP through BotGNNs results in better discriminators [4].

7 A good reason to consider dissimilar molecules is that it allows us to explore more diverse molecules.

8 It is likely that a BotGNN with access to the information in *BD* along with the Chemprop prediction would result in a better proxy model. We do not explore this here.

